# Flexible Adjustment of Oxytocin Neuron Activity in Mother Mice Revealed by Microendoscopy

**DOI:** 10.1101/2024.09.18.613777

**Authors:** Kasane Yaguchi, Kazunari Miyamichi, Gen-ichi Tasaka

**Affiliations:** Laboratory for Comparative Connectomics, RIKEN Center for Biosystems Dynamics Research, Kobe, Hyogo 650-0047, Japan; Graduate School of Biostudies, Kyoto University, Kyoto, Kyoto 606-8501, Japan; Japan Science and Technology Agency, PRESTO, Kawaguchi, Saitama 332-0012, Japan

**Author notes:** These authors contributed equally (Kasane Yaguchi, Gen-ichi Tasaka).

**Keywords:** Oxytocin, Milk ejection reflex, Microendoscopy, Paraventricular hypothalamus, Pup retrieval

## Abstract

Oxytocin (OT) neurons in the hypothalamic paraventricular nucleus (PVH) play an important role in a range of physiological and behavioral processes, including the initiation of milk ejection and the regulation of parental behaviors in mothers. However, their activity patterns at the single-cell level remain poorly understood. Using microendoscopic Ca^2+^ imaging in freely moving mother mice, we demonstrate highly correlated pulsatile activity among individual OT neurons during lactation. The number of OT neurons engaged in the pulsatile activity, along with the characteristics of individual waveforms, was dynamically modulated by lactation and weaning experiences. Notably, only ∼10% of the imaged OT neurons exhibited a significantly elevated response during pup retrieval, a hallmark of maternal behaviors, with a magnitude 18 times smaller than that observed during lactation. Collectively, these findings demonstrate the utility of microendoscopic imaging for PVH OT neurons and highlight the flexible adjustments of their individual activity patterns in freely behaving mother mice.

## INTRODUCTION

Oxytocin (OT), a nonapeptide hormone produced by OT neurons in the paraventricular (PVH) and supraoptic (SO) nuclei of the hypothalamus, plays an important role in facilitating uterine contractions during parturition, triggering milk ejection during lactation, and promoting maternal care toward infants [1–3]. PVH OT neurons also contribute to the reproductive functions of males [4], social bonding [5, 6], and the regulation of food intake and energy expenditure [7–9]. However, the way in which this small population of neurons can perform such a diverse array of functions remains an open question. Recording the activity of PVH OT neurons *in vivo* during behavioral and physiological processes would be an initial step toward understanding their functional heterogeneities. Previous research using single-unit *in vivo* electrophysiological recordings combined with optogenetic identification of OT neurons (opto-tag) demonstrated activity in these neurons when virgin female mice observed maternal behavior [10], when mother mice heard their pups’ ultrasonic vocalizations [11], and when female rats experienced social touch [12]. However, these previous studies are limited by their relatively small number of single units recorded from each animal. Two-photon Ca^2+^ imaging could allow spatiotemporal activity analyses of a larger number of OT neurons, albeit with the drawback of restricted mobility due to head fixation [13]. Overall, the activity patterns of PVH OT neurons during lactation and maternal behaviors have remained elusive.

The milk ejection reflex is initiated by suckling stimuli at the nipples, which subsequently trigger the pulsatile synchronous burst activities of PVH and SO OT neurons, leading to a transient increase in plasma OT. This OT pulse reaches the mammary gland, inducing the active transfer of milk from alveolar storage to mammary ducts [3]. Classical studies have most extensively characterized OT neuron activity through *in vivo* extracellular recording techniques in rats and rabbits, either under anesthesia [3, 14–16] or awake [17, 18]. These studies have identified the bursting activities of putative OT neurons during the milk ejection reflex, noting a ∼20–40-fold increase in the firing rate of individual neurons [14]. Paired-recording of PVH and SO OT neurons demonstrated that the peak of pulsatile activity was closely synchronized, with a mean time-lag of only ∼120 milliseconds. However, due to technical limitations, the identity of OT neurons in these studies was speculative, mostly based on their electrophysiological properties and axonal projection to the posterior pituitary, not on *OT* gene expression. Additionally, previous studies were limited by the small number of units that could be analyzed in each experiment in which long-term chronic recording was not feasible.

More recently, our group and others have visualized the pulsatile population activity of genetically defined PVH OT neurons of lactating mother mice using fiber photometry [19–21]. These studies have shown that i) pulsatile activity of PVH OT neurons exclusively occurs when multiple pups provide simultaneous suckling stimuli at the nipples; ii) the amplitude of this pulsatile activity increases, with the waveforms broadening, as mothers gain more lactation experience; and iii) this enhancement is intrinsic to the mother’s OT system, as fostering pups of different ages did not affect the peak height of photometry signals [19]. However, the spatial resolution of fiber photometry is insufficient to elucidate the underlying mechanisms at the single-cell level. Additionally, whether PVH OT neurons engaged in the milk ejection reflex are also active during maternal behaviors remains unknown.

Recent advancements in optical imaging tools have facilitated the visualization of individual neurons and the monitoring of their activities on a physiologically relevant time scale. For instance, head-mounted microendoscopes [22], combined with the Ca^2+^ sensor GCaMP [23], enable the visualization of activity in several dozen genetically defined neurons during natural behaviors. When paired with optical gradient refractive index (GRIN) lenses implanted in the brain, microendoscopes can target and image from deep brain regions such as the medial amygdala [24], ventromedial hypothalamus [25], and preoptic areas [26] within the context of social interactions. These innovations prompted us to apply a microendoscope to PVH OT neurons to characterize their activity patterns at single-cell resolution in freely behaving lactating mother mice.

## RESULTS

### Pulsatile activity dynamics of OT neurons across different postpartum days

We utilized a previously developed adeno-associated virus (AAV) to drive GCaMP6s expression under the control of the *OT minipromoter* (*OTp*), selectively targeting PVH OT neurons [20]. AAV9 *OTp-GCaMP6s* was injected into the PVH in the right hemisphere of adult female wild-type mice of the F1 hybrid of C57BL/6 and FVB strains, which shows robust maternal behaviors [27]. A GRIN lens (500-μm diameter) was implanted above the PVH at 3 weeks after the viral injection (Fig. 1A). GCaMP6s expression and the lens location were confirmed histologically (Fig. 1B and Fig. S1A). Figure 1C and 1D shows a representative example of a lactating mother on postpartum day (PPD) 6, with 37 regions of interest (ROIs; putatively corresponding to individual cells) within the field of view. After allowing the mother to interact freely with 8–10 pups in the home cage, we observed synchronous elevation of Ca^2+^ signals in all but one ROI (#30 in orange in panel D), with interpeak intervals of a few minutes, closely resembling the nature of pulsatile activity of PVH OT neurons previously characterized by fiber photometry during the milk ejection reflex (Movie S1) [19–21]. We defined the peak time as the moment when population-averaged activity reached its maximum for further analysis (Fig. 1E, F). A 30-second time window between 30 and 60 seconds after the peak was used as the baseline activity. Noise correlation analysis revealed a strong correlation between individual ROIs during pulsatile activity, with little or no correlation among baseline activities (Fig. 1G). These data confirmed the highly synchronous nature of PVH OT neurons, as previously suggested by a small number of single-unit electrophysiological recordings in rats [3, 14, 16], and demonstrated that microendoscopy is suitable for detecting the pulsatile activities of PVH OT neurons in mother mice.

**Figure 1.**
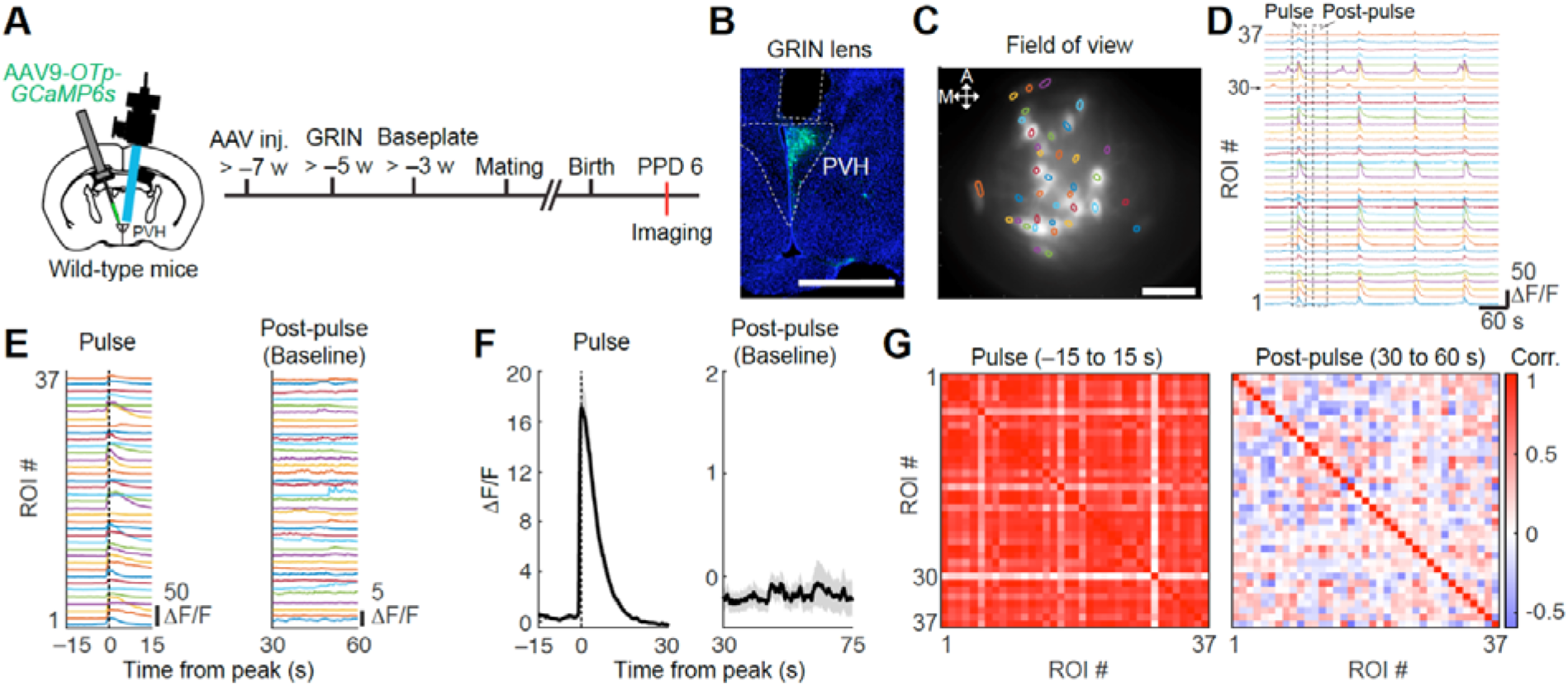
*In vivo* microendoscopic Ca^2+^ imaging of PVH OT neurons. (A) Schematic of the experimental time line. PPD, postpartum day. (B) Representative image showing the GRIN lens tract and the expression of *GCaMP6s* (green). Blue, DAPI counterstaining. Scale bar, 1 mm. (C) Example of a spatial map showing ROIs with the maximum projection image in a mother at PPD 6. Scale bar, 100 μm. (D) Examples of Ca^2+^ responses from ROIs during lactation. The indicated colors correspond to those in panel C. (E) Representative Ca^2+^ responses from individual ROIs during and after the pulsatile event indicated by the dotted line in panel D. The peak time of population-averaged pulsatile activity was set to 0. (F) Population-averaged activity during the pulsatile event marked by the dotted line in panel D. Shadows represent SEM. (G) Noise correlation matrices between individual ROIs during pulsatile and post-pulsatile activities. Among these ROIs, only ROI#30 did not exhibit synchronized pulsatile activity.

To recapitulate an increase in pulsatile population activities of PVH OT neurons from the early (PPD 1–2) to middle (PPD 11–12) lactation stages [19], we first collected microendoscopic data from mothers at PPD 1–2, PPD 5–6, and PPD 11–12 (Fig. 2A, Movie S1) and measured the photometry-like signals by extracting Ca^2+^ transients from a single large ROI covering the entire GRIN lens (Fig. S1B, C). We found a larger peak amplitude and a broader waveform, as defined by the full-width at half maximum (FWHM), on PPD 11–12 compared with PPD 1–2 (Fig. S1D–F), confirming previous photometry data [19].

**Figure 2.**
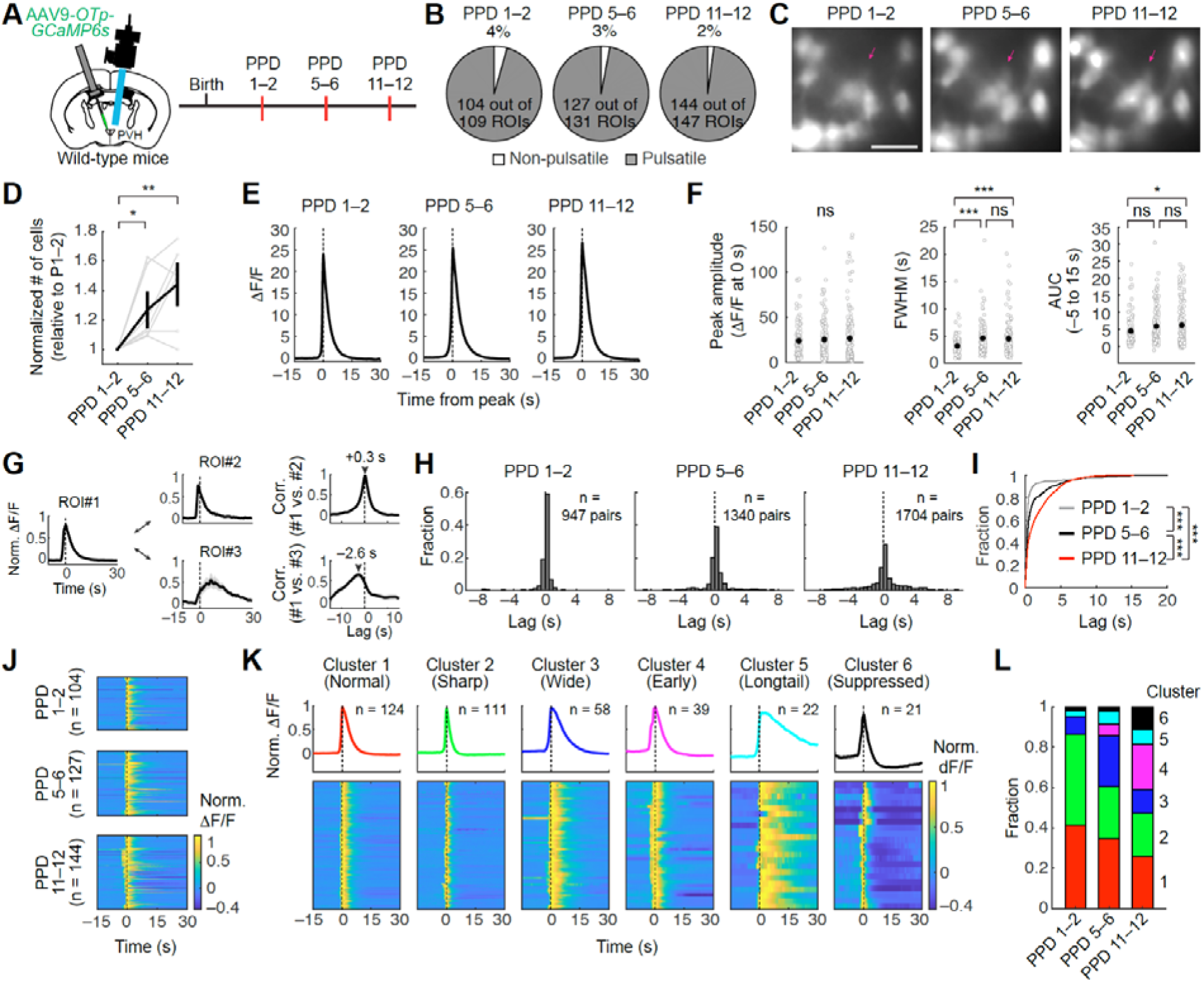
Dynamics of pulsatile activity in OT neurons across different postpartum days. (A) Schematic of the experimental time line. (B) Fraction of ROIs showing pulsatile activity across different lactation stages. Pooled data from n = 7 mice. (C) Representative maximum projection images during pulsatile activity (over 30 seconds) from different lactation stages. Arrows indicate the ROIs detected at PPD 5–6 and PPD 11–12, but not at PPD 1–2. Scale bar, 50 μm. (D) Number of ROIs showing pulsatile activity for each mouse. N = 7 mice. Error bars, SEM. (E) Averaged activity traces during the pulsatile event. The peak time for each ROI was set to time 0. (F) Quantification of the peak amplitude (left), FWHM (middle), and AUC between –5 and +15 seconds (right). *, p < 0.05 and ***, p < 0.001 by post hoc Tukey’s HSD test after a significant one-way ANOVA test. (G) Cross-correlation analysis examples from three ROIs that were simultaneously recorded. Normalized ΔF/F traces aligned to the population-averaged peak at time 0 (left). Cross-correlation results between ROI#1 and ROI#2 (top right) and between ROI#1 and ROI#3 (bottom right). The peak of the correlation coefficient is indicated with an arrowhead and corresponding time lags. (H) Distribution of time lags from cross-correlation analysis across three different lactation stages. Histograms show the fraction of pairs with specific time lags for each group. Indicated number is the number of pairs. (I) Cumulative plots of the distribution of peak time lag from different lactation stages. ***, p < 0.001 by the Kolmogorov–Smirnov test with Bonferroni correction. (J) Heat map representations of normalized and averaged responses of individual ROIs aligned to the population-averaged peak at time 0. (K) Classification of ROIs into six clusters based on their event-averaged activity (see Methods and Fig. S2). Upper traces represent the population-averaged activity during pulsatile activity within each cluster, and bottom heat maps represent event-averaged responses of individual ROIs from each cluster. (L) Fractions of ROIs classified into each cluster across different lactation stages. Shadows (in panels E and G) represent SEM.

Further analyses indicated that over 96% of recorded ROIs from a total of seven mothers exhibited pulsatile activity (Fig. 2A, B). The remaining few percent of neurons may represent nonspecifically labeled non-OT neurons, given that the targeting specificity of *OTp* is approximately 95% [20]. The number of ROIs with significant responses during pulsatile activity gradually increased over the postpartum period (Fig. 2B, C), resulting in a 1.4-fold increase in responding ROIs at PPD 11–12 compared with PPD 1–2 (Fig. 2D). The response magnitude of individual ROIs, determined by the peak amplitude of peri-event time histograms of pulsatile activity (Fig. 2E), did not differ among groups (Fig. 2F, left). By contrast, the average FWHM gradually and significantly increased (Fig. 2F, middle). Consequently, the area under the curve (AUC) of the pulsatile activity significantly increased at PPD 11–12 (Fig. 2F, right). These results suggest that more PVH OT neurons are recruited to pulsatile activity by the middle lactation period, with cells displaying broader pulsatile activity, leading to larger and wider pulsatile activity at the population level, as observed in previous studies using fiber photometry recordings [19, 21]. These changes may contribute to the larger plasma OT pulse in the later stage of lactation, when the mother needs to provide more milk to the infants than in the early stage of lactation, as demonstrated in rabbits [28].

To characterize the temporal dynamics of pulsatile activity among individual cells, we next compared the cross-correlation for each pair of ROIs concurrently recorded during lactation. Cross-correlation analysis quantifies the temporal relationship between the activity patterns of two neurons, revealing the degree of synchronization and potential time lags [29, 30]. Fig. 2G shows examples of cross-correlation analysis from three ROIs that were recorded simultaneously at PPD 6. We analyzed the distribution of time lags from cross-correlation analyses at different lactation stages (Fig. 2H). The time lags for most pairs were small, with 88.1% of total pairs being within a 1-second time lag at PPD 1–2, indicating a high degree of synchronicity in pulsatile activity during lactation. By contrast, later lactation stages exhibited a broader cumulative distribution curve of the time lag, suggesting increased temporal variability with advances in lactation stage (Fig. 2H and 2I).

Next, to evaluate the heterogeneity in pulsatile activity patterns across different lactation stages, we performed an unbiased clustering analysis on the event-averaged responses during pulsatile activity (Fig. 2J and Fig. S2, also see Methods). We identified six clusters of ROIs based on their activity during pulse events (Fig. 2K and Fig. S2D). We qualitatively annotated these clusters as follows: normal (Cluster 1 - Normal), sharp pulse form (Cluster 2 - Sharp), wide pulse form (Cluster 3 - Wide), early onset (Cluster 4 - Early), sustained longtail form (Cluster 5 - Longtail), and post-suppressed activity pattern (Cluster 6 - Suppressed). The majority of ROIs at PPD 1–2 belonged to Clusters 1 and 2, which represented relatively stereotypical pulsatile activity patterns, comprising 87% of the population. By contrast, Clusters 3–6 expanded in the middle lactation stages compared with PPD 1–2, while the relative proportion of Clusters 1 and 2 decreased (Fig. 2L and Fig. S2E). Particularly, Clusters 4–6 covered up to 41% of the population at PPD 11–12. These data suggest that the pulsatile activity patterns of individual PVH OT neurons can be dynamically modulated from the early to middle lactation stages.

### Tracking of the same OT neurons across different postpartum days

By far, our data revealed two populations of PVH OT neurons: one group engaged in pulsatile activity from PPD 1, and the other newly recruited as lactation progressed. To explore the physiological differences between these populations, we tracked the same ROIs across different PPDs, taking advantage of the sparse labeling of OT neurons (Fig. S3). We focused on PPD 1–2 and PPD 11–12, as differences became more pronounced at PPD 11–12 than at PPD 5–6 (Fig. 2I). We define ROIs persisting across days as “Maintained”, those recruited in later lactation as “Recruited”, and those not detected later as “Disappeared” (Fig. 3A). At PPD 11–12, 97 of 144 ROIs were classified as “Maintained”, and 47 as “Recruited” (Fig. 3B). Only seven ROIs were classified as “Disappeared”.

**Figure 3.**
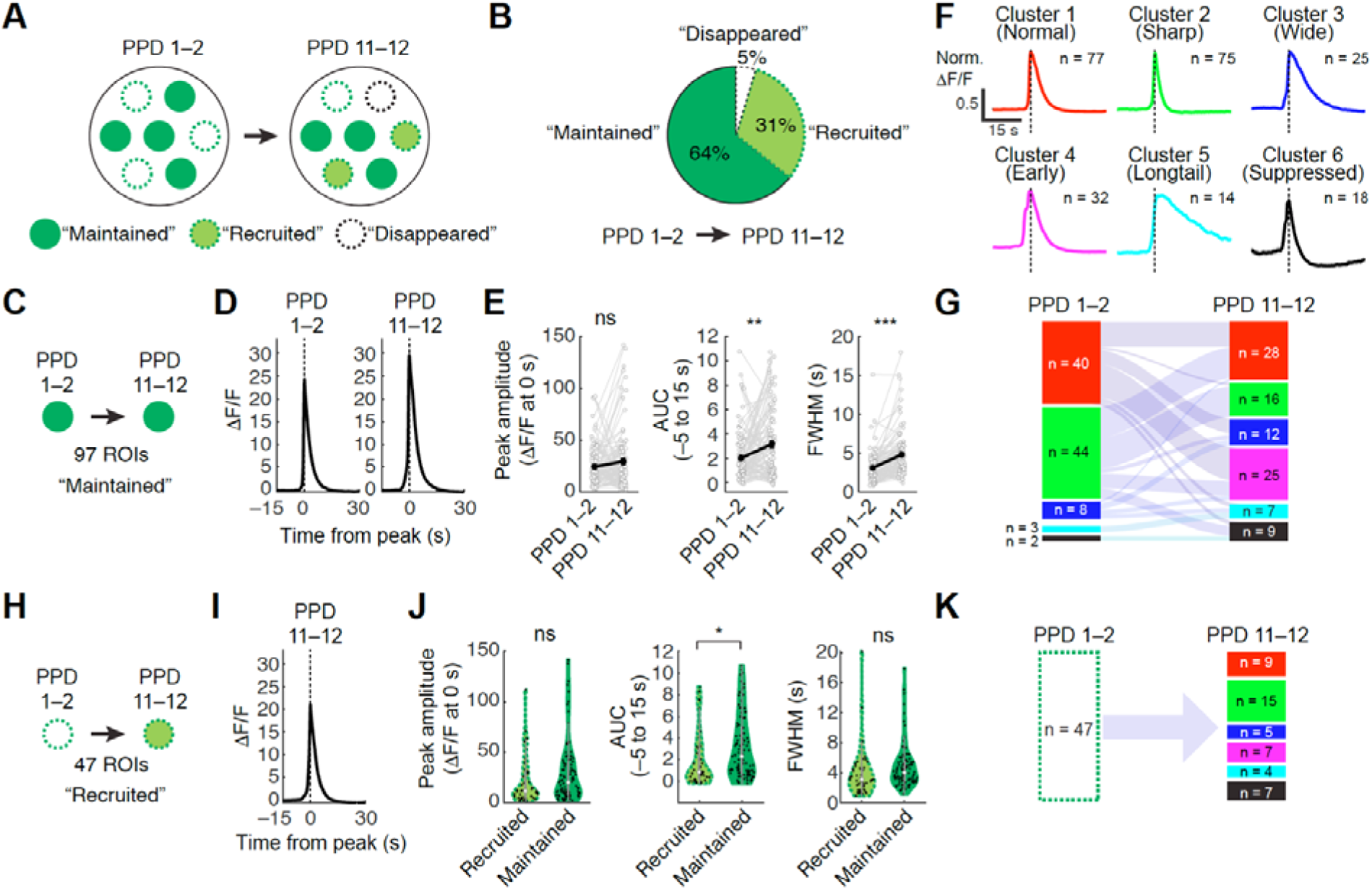
Longitudinal tracking across days of imaging. (A) Schematic of three distinct types of OT neurons showing pulsatile activity. (B) Fraction of ROIs belonging to the three different types. Pooled data from n = 7 mice. (C) Schematic of the “Maintained” cells. (D) Averaged activity traces during pulsatile activity from ROIs tracked successfully between PPD 1–2 and PPD 11–12. The peak time for each ROI was aligned to 0. (E) Quantification of the peak amplitude (left), AUC between –5 and +15 seconds (middle), and FWHM (right). *, p < 0.05 and ***, p < 0.001 by post hoc HSD test after a significant one-way ANOVA test. (F) Population-averaged activity during pulsatile activity within each cluster. (G) Sankey diagram showing the retention or transition of clusters within the “Maintained” population between different lactation stages. (H) Schematic of the “Recruited” cells. (I) Averaged activity traces during pulsatile activity. (J) Quantification of the peak amplitude (left), AUC between –5 and +15 seconds (middle), and FWHM (right). *, p < 0.05 by the Mann–Whitney *U* test. (K) Number and fraction of assigned clusters within the “Recruited” population. Shadows (in panels D and I) represent SEM.

We first analyzed the “Maintained” ROIs (Fig. 3C). These ROIs exhibited a significant increase in FWHM and AUC of pulsatile activity, mirroring trends observed in the population analysis shown in Fig. 2 (Fig. 3D, E). We then examined the cluster transitions based on pulsatile activity patterns (Fig. 3F, G). Many ROIs from Clusters 1 and 2 at PPD 1–2 displayed broader pulse forms at PPD 11–12 (Fig. 3G). For example, 15 of 40 Cluster 1 ROIs at PPD 1–2 transitioned to Clusters 3–6 at PPD 11–12. By contrast, ROIs within Clusters 5 and 6 at PPD 1–2 remained stable. Notably, nearly all ROIs in Cluster 2 at PPD 11–12 originated from Cluster 2 at PPD 1–2, indicating that almost no cells exhibited a narrowing of the pulse shape. We next analyzed “Recruited” ROIs (Fig. 3H). These ROIs showed a significantly smaller AUC compared with the “Maintained” ROIs, with no significant difference in peak amplitude and FWHM (Fig. 3I, J). Among “Recruited” ROIs, all clusters were present, with no biased distribution from the fraction observed in the entire population (Fig. 3K, compared with Fig. 2L).

We recognize a few technical limitations of our imaging system: i) given the low baseline activity observed in lactating mothers, some PVH OT neurons may be undetectable unless they display substantial pulsatile activity; and ii) variability in cell detection may arise because of the limited z-axis resolution, even for neurons with consistent pulsatile activity. Therefore, we cannot fully rule out the possibility that “Recruited” cells might exhibit weak activity below the detection threshold or be located slightly out of focus at PPD 1. Despite these limitations (see Discussion), our data demonstrated a significant increase in the intensity of pulsatile activity in “Recruited” cells and substantial cluster transitions, leading to a broadening of the waveform in “Maintained” cells, as lactation progresses.

### A subset of PVH OT neurons exhibits a time-locked response to pup retrieval

We next performed microendoscopic Ca^2+^ imaging during pup retrieval, followed by lactation, to compare the responses of PVH OT neurons during these episodes (Fig. 4A). For pup retrieval, we conducted a 6-minute recording session where new pups were promptly presented upon the retrieval of existing ones (Fig. 4B). Pup retrieval comprises three major behavioral categories in the following sequence: contact with a pup, onset of retrieval, and dropping a pup into the nest (completion) (Fig. 4C) [31]. We observed that certain ROIs exhibited an increased Ca^2+^ transient (referred to as an elevated response) in response to the onset of retrieval (Fig. 4D). We recorded a total of 127 ROIs from seven mice and found that 10% of ROIs exhibited increased and another 10% exhibited decreased Ca^2+^ transients (referred to as elevated or suppressed responses, respectively) synchronized with one of the three behavioral categories (Fig. 4E–H). The majority of responses were time-locked to the onset of retrieval (Fig. 4F).

**Figure 4.**
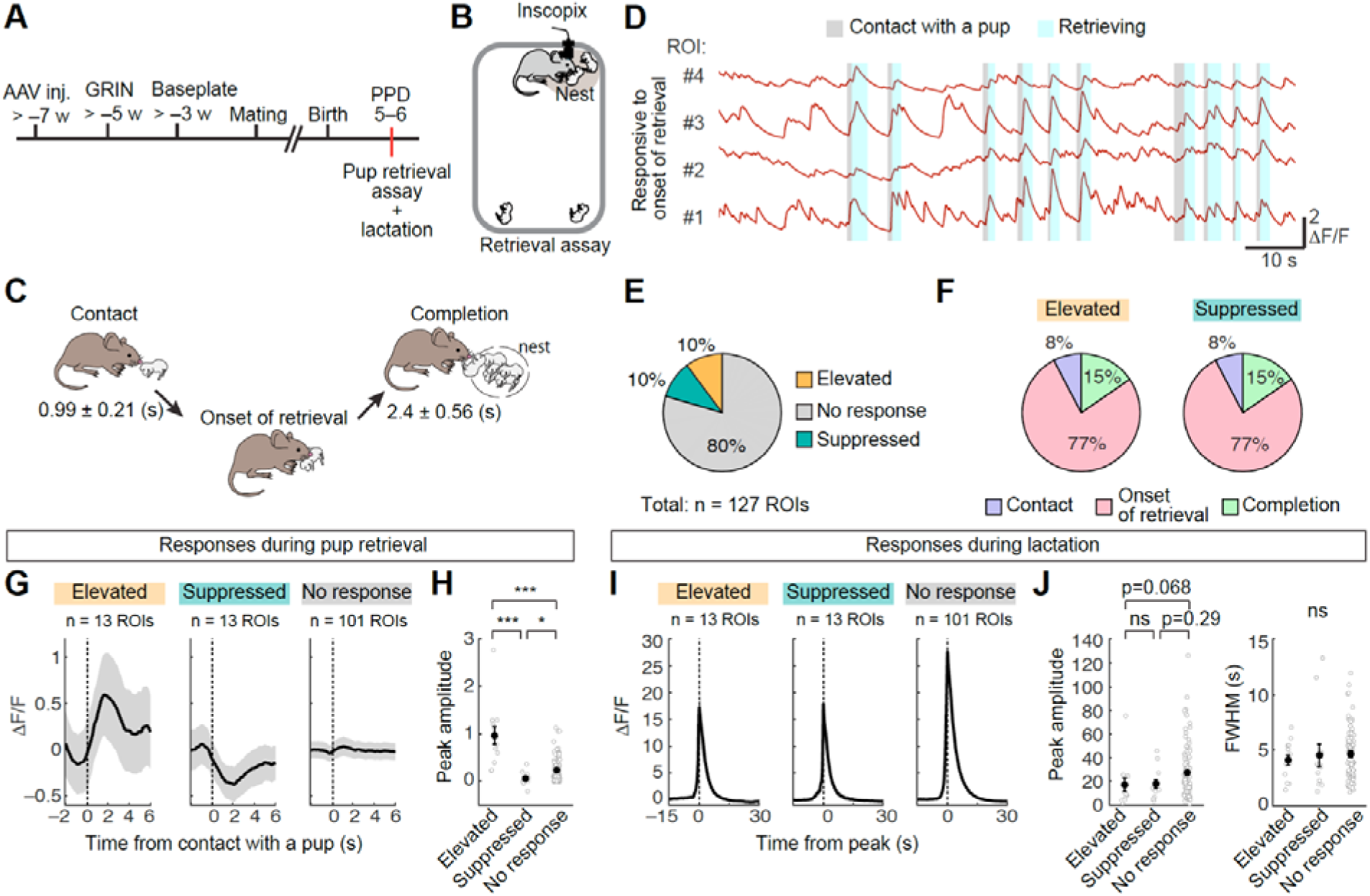
Responses of PVH OT neurons during pup retrieval. (A) Schematic of the experimental time line. (B) Schematic of the pup retrieval assay during Ca^2+^ imaging. (C) Schematic of sequential behavioral categories during pup retrieval. The number indicates the duration between events (± standard deviation). (D) Examples of Ca^2+^ responses from four ROIs responding to onset of retrieval within a single animal. (E) Fraction of elevated (orange) and suppressed (blue) responses during pup retrieval. (F) Fraction of ROIs that responded to one of the three behavioral categories of pup retrieval among the elevated (left) or suppressed (right) population. (G) Averaged activity traces of significantly responsive and nonresponsive ROIs to the contact with a pup. (H) Quantification of mean peak amplitude. *, p < 0.05 and ***, p < 0.001 by post hoc Tukey’s HSD test after a significant Kruskal-Wallis test. (I) Averaged activity traces during the pulsatile event. Peak time for each ROI was set to 0. (J) Quantification of mean peak amplitude (left) and FWHM (right). Indicated p-values were measured by post hoc Tukey’s HSD test after a significant Kruskal-Wallis test. Error bars, SEM. Shadows (in panels G and I) represent SEM.

We also explored the activity of these retrieval-responsive OT neurons during lactation. ROIs with both elevated and suppressed responses during pup retrieval showed significantly increased activity during lactation, with the magnitude of this response being approximately 18-fold greater during lactation than that observed during pup retrieval (compare Fig. 4G and I in elevated population). Additionally, we observed a slightly weaker trend of lactation-related pulsatile activity in OT neurons with elevated responses during pup retrieval compared with those with suppressed or no responses (Fig. 4J). These data show the heterogeneity of PVH OT neurons and highlight the utility of microendoscopic imaging in detecting relatively weak activity of PVH OT neurons during pup retrieval.

### The pulsatile activity of OT neurons during the post-weaning period in the absence of milk letdown

While the pulsatile secretion of OT is essential for milk ejection, it remains unclear whether this activity in PVH OT neurons is exclusive to postpartum lactating mothers. Human studies suggest that women can reestablish lactation after having stopped for months, a process known as “relactation” [32–34]. To test whether the pulsatile activity of PVH OT neurons occurs in post-weaning mothers following suckling stimuli by pups, we conducted photometry recordings from post-weaning female mice who underwent regular parturition and lactation until around PPD 21, when their pups were weaned. At 2 weeks after weaning, following confirmation of their restored estrus cycles via vaginal cytology [35], these female mice were co-housed with a lactating mother and pups for 1 day for habituation. At around PPD 37, the post-weaning mother mice were co-housed with novel 12-day-old foster pups without lactating mothers for fiber photometry recording (Fig. 5A). We observed pulsatile activity of PVH OT neurons (Fig. 5B) that closely resembled that observed at PPD 12 in the same mice. However, the body weight of the pups was significantly reduced after the imaging session, indicating insufficient milk letdown in the post-weaning mothers (Fig. 5C). A closer examination revealed that the frequency of pulsatile activity was significantly lower and the amplitude of each pulsatile activity was smaller during re-suckling at PPD 36–38 compared to those observed at PPD 11–12 during actual lactation (Fig. 5D–F). Additionally, the waveform of pulsatile activity was significantly shorter in post-weaning mother mice (Fig. 5F, right). These data indicate that i) the pulsatile activity of PVH OT neurons following suckling stimuli can be observed even after the reproductive cycle is restored in post-weaning mothers, and ii) the frequency, intensity, and waveform of this activity differ substantially between the postpartum lactation period and post-weaning re-suckling trials.

**Figure 5.**
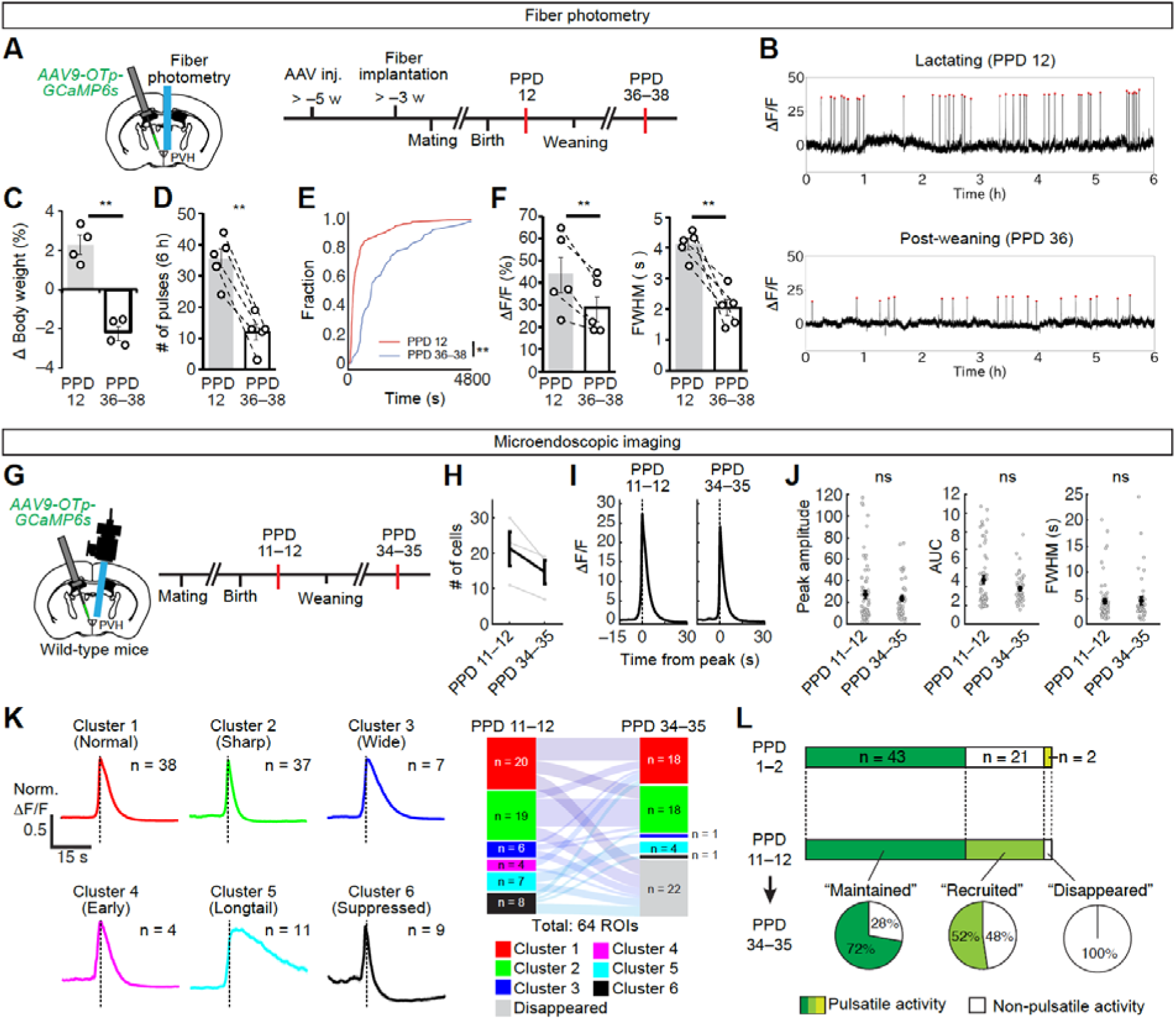
Pulsatile activity of PVH OT neurons during the post-weaning period. (A) Schematic of the experimental time line for fiber photometry. (B) Typical photometry traces on PPD 12 during lactation (top) and PPD 36 post-weaning (bottom). Red dots denote pulsatile activities of OT neurons. Data were recorded for 6 hours during the light phase. (C) Quantification of pup body weight changes after 6 hours of recording. **, p < 0.01 by an unpaired *t*-test. (D) Quantification of the number of pulsatile activities of OT neurons on PPD 12 and PPD 36–38. N = 5 mice. **, p < 0.01 by a paired *t*-test. (E) Cumulative distribution of inter-pulse intervals comparing patterns on PPD 12 (red) and PPD 36–38 (blue). **, p < 0.01 by the Kolmogorov–Smirnov test. (F) Quantification of the peak amplitude (left) and FWHM (right). N = 5 mice. **, p < 0.01 by a paired *t*-test. (G) Schematic of the experimental time line for microendoscopic imaging. (H) Number of ROIs showing pulsatile activity for each mouse. N = 3 mice. (I) Averaged activity traces during the pulsatile event. Peak time for each ROI was aligned to 0. Shadows represent SEM. (J) Quantification of the mean peak amplitude (left), AUC between –5 and +15 seconds (middle), and FWHM (right). **, p < 0.01 and ***, p < 0.001 by post hoc Tukey’s HSD test after a significant one-way ANOVA test. (K) Left, traces represent population-averaged activity during pulsatile activity within each cluster. Right, Sankey diagram showing the retention or transition of clusters in the population exhibiting pulsatile activity on PPD 11–12. (L) Fraction of ROIs classified as “Maintained”, ”Recruited”, and “Disappeared” based on their response patterns from PPD 1–2 to PPD 11–12 (upper). The bottom pie charts represent the fractions exhibiting or not exhibiting pulsatile activity on PPD 34–35. Error bars, SEM.

To elucidate these alterations in photometry signals at single-cell resolution, we employed microendoscopic imaging while mother mice cared for their pups at PPD 11–12, and when the same females were exposed to foster pups at 2 weeks post-weaning at PPD 34–35 (Fig. 5G). We used 2-day-old foster pups instead of 12-day-old ones to observe the milk tank directly. Consistent with the fiber photometry data, the pup’s body weight declined following the imaging session (Fig. S4A) with empty milk tanks, suggesting inadequate milk letdown in the post-weaning mothers. Nevertheless, pulsatile activity of PVH OT neurons was observed. Measurements of Ca^2+^ transients from a single large ROI covering the entire GRIN lens confirmed an overall reduction in the intensity of population pulsatile activity from PPD 11–12 (actual lactation) to PPD 34–35 (post-weaning re-suckling) (Fig. S4B, C), as well as a decrease in waveform width (Fig. S4D). Analysis of individual ROIs revealed a reduction in the number of responsive ROIs at PPD 34–35 compared with PPD 11–12 (Fig. 5H). Basic properties of pulsatile activity, such as the peak amplitude and FWHM during pulsatile activity, showed no significant differences (Fig. 5I, J).

To evaluate the pulsatile activity pattern, we tracked the same ROIs from PPD 11–12 (actual lactation) to PPD 34–35 and assigned the data from PPD 34–35 to the existing six clusters in lactating mothers used in Fig. 2G (Fig. 5K, see Methods). Overall, the number of ROIs in Clusters 1 and 2 remained mostly unchanged as a result of relatively stable ROIs staying in the same cluster from PPD 11–12 to PPD 34–35, in addition to ROIs transitioning from wider clusters (Clusters 3–5) to Cluster 1. By sharp contrast, most ROIs in Clusters 3–6 at PPD 11–12 disappeared at PPD 34–35. These changes collectively make the proportion of clusters in post-weaning mothers similar to the early lactation stage, PPD 1–2, with Clusters 1 and 2 occupying the majority and a small fraction of Clusters 3–6. Finally, we explored the attribution of cells showing pulsatile activity at PPD 34–35 by longitudinal tracking analysis across PPD 1–2, PPD 11–12, and PPD 35–36. We found that 72% of “Maintained” ROIs at PPD 11–12 were retained at PPD 34–35, while 48% of “Recruited” ROIs disappeared at PPD 34–35 (Fig. 5L). Thus, “Recruited” cells at PPD 11–12 are more likely to return to a non-responding state after weaning, suggesting that they act as adjusters of pulsatile activity of PVH OT neurons across different states.

## DISCUSSION

Deciphering how distinct hormone-producing neurons in the hypothalamus orchestrate diverse physiological and behavioral processes represents a major challenge in neuroscience [36]. The present study utilized microendoscopic imaging to observe the suckling-induced pulsatile activity of PVH OT neurons at single-cell resolution in freely moving lactating and post-weaning mother mice. Our data highlight the highly dynamic nature of individual PVH OT neurons across lactation stages, with a small subset responding during pup retrieval. Here, we discuss the biological insights obtained from this study, along with limitations.

While classical extracellular recording studies have identified the bursting activities of putative OT neurons during the milk ejection reflex [3, 14, 15], the temporal dynamics of individual OT neuron activity across different lactation stages have remained elusive. Our data illuminate a heightened activity intensity of the “Recruited” population and broadening waveforms of the “Maintained” population from the early to mid-lactation stages (Figs. 2 and 3 and Fig. S5). The molecular and cellular characteristics of the “Maintained” and “Recruited” populations remain unclear and thus constitute an important subject for future investigations. Our unbiased clustering supports overall broadening changes from the early to mid-lactation stages, with broader clusters expanding at PPD 11–12. Tracking individual ROIs further revealed complex and dynamic changes of pulse waveforms across lactation stages. In addition, the synchronicity among PVH OT neurons decreased (Fig. 2G–I). These changes collectively contribute to the wider and more intensive population activity of PVH OT neurons. Assuming that OT release from the posterior pituitary reflects the integral of the population activity waveform, these changes may facilitate adjustments in the intensity and duration of milk ejection, ensuring adequate milk release from the mammary gland.

Although relactation is well-documented in humans [32–34], it was unclear whether PVH OT neurons in female mice could respond to suckling stimuli and exhibit pulsatile activity beyond the regular lactation period. Our data reveal that pulsatile activity can be reinstated at 2 weeks post-weaning. As insufficient or no milk letdown was observed during the re-suckling trials, the pulsatile activity of PVH OT neurons appears to be separable from lactogenesis, which is predominantly regulated by prolactin [37, 38]. Although the precise mechanism by which multiple OT neurons form synchronous pulsatile activity remains unclear, our data suggest that such activity patterns are not exclusive to a specific lactation period immediately following parturition. This view is consistent with classical studies suggesting that *exo-vivo* cultures of neonatal rat PVH neurons could exhibit synchronous pulsatile activity of putative OT neurons [39].

During these re-suckling trials, many pulsatile activity properties revert to a state similar to PPD 1: i) the inter-pulse interval elongates as observed at PPD 1 [20], ii) the number of significantly responsive neurons decreases, and iii) the waveform patterns return to a simpler state where Clusters 1 and 2 dominate. Points ii) and iii) collectively explain the narrower and smaller population activity seen in the post-weaning mothers, resembling those observed at PPD 1. Although the extent to which these changes are intrinsic to the mother’s OT system remains unclear, these data suggest that the post-weaning period may serve to reset the heightened pulsatile activity observed in mid-lactation, preparing for the next round of parturition and lactation. Consistent with this view, our previous study demonstrated that the population pulsatile activity of PVH OT neurons during the second round of lactation following weaning returns to levels comparable to those observed at PPD 1 in primiparous mother mice [19].

In addition to characterizing the suckling-induced pulsatile activity of PVH OT neurons, this study revealed an active subpopulation of PVH OT neurons during pup retrieval. While a previous study demonstrated similar pup retrieval-locked activity using fiber photometry in biparental male mandarin voles [40], our previous attempts failed to detect photometry signals during pup retrieval in lactating mother mice [20]. Based on our new data, this negative result can be explained by the following three reasons: 1) the neurons responding positively during pup retrieval constitute only approximately 10% of the imaged PVH OT neurons; 2) the activity intensity is less than 18-fold compared with suckling-induced pulsatile activity; and 3) a similar number of PVH OT neurons show negative (suppressed) responses during pup retrieval, potentially canceling the positive response at the population level. These response characteristics highlight the power of microendoscopy for single-cell resolution imaging. Furthermore, our data show that the suckling-induced pulsatile activity of the pup-retrieval-responding population is relatively weaker, suggesting that they may belong to a distinct subtype of PVH OT neurons. One intriguing possibility is that this pup retrieval-responding population belongs to parvocellular PVH OT neurons, a subtype that selectively projects to central brain regions, including the ventral tegmental area and substantia nigra, thereby modulating the dopamine system and contributing to reward processing [6, 41–43]. However, whether parvocellular PVH OT neurons precipitate in pulsatile activity during milk ejection remains unknown, and is therefore an important subject for future research.

While microendoscopic imaging of PVH OT neurons provides novel insights, as outlined above, we acknowledge four major limitations of the present method. First, the spatial resolution, particularly along the z-axis, is low in single-photon imaging. Fine dendritic and axonal processes of neighboring cells and cell bodies beneath the imaging plane may contaminate individual ROIs, compromising specificity. Second, due to the low basal activity of PVH OT neurons in mother mice, the identification of responsible ROIs primarily relies on the pulsatile activity, compromising the characterization of PVH OT neurons that do not exhibit pulsatile activity. Third, the number of ROIs per animal was as low as 20–30, much smaller than those obtained in the neocortex or hippocampus [44]. Fourth, the relatively low temporal resolution of GCaMP6s [23] precluded correlating the observed intensity of pulsatile activity with the number of action potentials. During the milk ejection reflex, individual PVH OT neurons can fire at rates up to ∼80 Hz [14]. Ideally, future studies should utilize miniaturized two-photon microscopy technology [45] with improved sensors to overcome these limitations. We anticipate that *in vivo* imaging of hormone-producing PVH neurons in freely behaving animals will continue to enhance our understanding regarding the hypothalamic control of behavioral and physiological processes.

## Acknowledgments

We thank Takuya Osakada (UY University), Tomomi Karigo (Johns Hopkins University), and the members of the Miyamichi lab for the critical reading of the manuscript, Ayumu Konno and Hirokazu Hirai (Viral Vector Core, Gunma University Initiative for Advanced Research) for AAV *Otp-GCaMP6s*, and the RIKEN BDR animal facility for taking care of the animals. This work was supported by the RIKEN Junior Research Associate Program to K.Y., the JST PRESTO program (JPMJPR21S7), JSPS KAKENHI (20K15941), and RIKEN incentive and diversity promotion grants to G.T., and JSPS KAKENHI (21H02587 and 23H04945) and RIKEN BDR Stage Transition Project grants to K.M.

## Author Contributions

K.Y., K.M., and G.T. conceived the experiments. K.Y. performed the photometry recording experiments and analyzed the data. G.T. performed the endoscopic imaging experiments and analyzed the data. K.Y., K.M., and G.T. wrote the paper.

## Materials and Methods

### Animals

All animals were housed under a regular 12-hour light/dark cycle with *ad libitum* access to food and water. Wild-type FVB mice were purchased from CLEA Japan, Inc. (Tokyo, Japan) to generate the F1 hybrid of C57BL/6 and FVB strains for the microendoscopic imaging experiments (age range, 2–5 months). Wild-type C57BL/6 female mice (age range, 2–5 months) were used for the fiber photometry recording experiments. All experimental procedures were approved by the Institutional Animal Care and Use Committee of the RIKEN Kobe branch. Given that our study focused on lactation and maternal behaviors, we exclusively utilized female mice.

### Viral preparations

*pAAV-Otp-GCaMP6s* (Addgene #192945) was constructed as previously described [20]. The AAV serotype 9 *Otp-GCaMP6s* was created at the Gunma University Viral Vector Core.

### Stereotaxic injection

For targeting AAV into specific brain regions, stereotaxic coordinates were first defined for each brain region based on the Allen Brain Atlas [46]. Mice were anesthetized with 65 mg/kg ketamine (Daiichi-Sankyo) and 13 mg/kg xylazine (Sigma-Aldrich) via intraperitoneal injection and then head-fixed to stereotaxic equipment (RWD or Narishige). To image the neural activity of OT neurons in the PVH, 200 nL of AAV9 *Otp-GCaMP6s* (titer: 2.9 × 10^12^ genome particles per mL) was injected into the bilateral PVH (coordinates: 0.5 mm posterior, 0.2 mm lateral from the bregma, and 4.4 mm ventral from the brain surface).

### *In vivo* microendoscopic imaging

For microendoscopic recording, a ProView GRIN lens (500-μm diameter, 8.4 mm length, Inscopix) was inserted into the PVH (coordinates: 0.5 mm posterior, 0.2 mm lateral from the bregma, and 4.4 mm ventral from the brain surface with a 5° tilt from vertical). Mice were anesthetized with 65 mg/kg ketamine (Daiichi-Sankyo) and 13 mg/kg xylazine (Sigma-Aldrich) via intraperitoneal injection and head-fixed to stereotaxic equipment (Narishige). A 1-mm diameter craniotomy was performed over the lens target area, removing any remaining bone and overlying dura using fine forceps. Brain tissue was aspirated to a depth of 3 mm from the surface. A GRIN lens was mounted onto the ProView lens holder and attached to the Inscopix nVista. This unit was slowly lowered into the brain while monitoring *GCaMP6s* expression through the nVista. Upon reaching the desired depth, the lens was permanently fixed using Super-Bond (Sun Medical) and sealed to the skull. A metal bar (CF-10; Narishige) was also affixed to facilitate easy attachment and detachment of the microscope. After the adhesive had fully set, the camera and lens holder were carefully detached from the lens, and Kwik-Kast (WPI) was used to protect the exposed lens surface. After more than 3 weeks of recovery, the mice were anesthetized and placed in the stereotaxic equipment again. The focal plane was adjusted to bring *GCaMP6s*-labeled cells into focus, and a baseplate (Inscopix) was permanently glued with Super-Bond. After more than 1 week of recovery, the microscope was attached, and the mice were allowed to explore freely in their home cage for 5–10 minutes. This habituation session was conducted more than twice before the first imaging session.

We performed microendoscopic imaging using the Inscopix nVista system (Inscopix) without refocusing across imaging sessions within 1 day, while the focal plane was adjusted each day before the first imaging session. Before imaging, the microendoscope was attached to the animals by securing the implanted head bar. Images (1080 × 1080 pixels) were acquired using nVista HD software (Inscopix) at 10 Hz, with LED power set to 0.4–0.8 mW/mm^2^ and a gain of 2.0–3.0. Timestamps of the imaging frames and camera data were collected for alignment using WaveSurfer (https://wavesurfer.janelia.org/). Several imaging sessions were conducted, each lasting for 15 minutes, with a few minutes’ interval between sessions. The imaging data were cropped to 800 × 700 pixels and exported as .tiff files using the Inscopix data processing software. To identify putative cell bodies for neural signal extraction, we used v2 of MIN1PIPE (https://github.com/JinghaoLu/MIN1PIPE [47]) with a spatial down-sampling rate of 2. All traces from identified cells were manually inspected to ensure signal quality, and excluded if they exhibited abnormal shapes or overlapped signals from adjacent cells. Relative changes in calcium fluorescence were calculated by ΔF / F_0_ = (F – F_0_) / F_0_ (where F_0_ represents the median fluorescence of the entire trace). For longitudinal tracking, the same cells across different days were manually detected.

### Measurement of photometry-like signals from the microendoscopic imaging data

To quantify photometry-like signals from the microendoscopic imaging data, a large ROI covering the entire GRIN lens was drawn, and the average signal for each frame was calculated using custom-written MATLAB code. For background signal calculation, the average signal from an ROI outside the GRIN lens was measured. A fitted signal was generated by applying least-squares linear fitting to align this background signal with the *GCaMP6s* signal using the MATLAB *polyfit* function. ΔF/F was obtained by subtracting the fitted background signal from the *GCaMP6s* signal to eliminate movement and other common artifacts.

### Detection of pulsatile activity

To detect pulsatile activity events, mean traces of ΔF/F from all simultaneously recorded ROIs were calculated. Peaks were then identified using the MATLAB *findpeaks* function, with peak times defined as time 0. To assess whether each ROI showed pulsatile activity, noise correlation and cross-correlation between an ROI and the population-averaged pulsatile activity were calculated using a time window ranging from –15 to 15 seconds relative to the peak time of the population-averaged pulse event. ROIs were classified as exhibiting pulsatile activity if they had a noise correlation above 0.25 and a cross-correlation above 0.65. The values and lags of the cross-correlation between a pair of ROIs were calculated using the mean cross-correlation taken for each of the multiple pulses using MATLAB *xcorr* function.

To determine the pulse timing for individual cells, a time window ranging from –15 to 15 seconds from the peak of the population-averaged pulse event was used, and the peaks for each ROI were detected using the MATLAB *findpeaks* function, with a peak height threshold of 0.1. Due to varying peak heights across cells, ΔF/F traces were normalized so that the peak of the entire trace was set to 1. If no peak was detected under these conditions, the peak timing of the population-averaged pulsatile activity was used as time 0 for that particular pulsatile activity event.

### Clustering analysis

We performed clustering analysis on the averaged responses during pulsatile activity to identify potential functional clusters and evaluate the heterogeneity in pulsatile activity patterns. Averaged pulsatile activity data ranging from –15 to 30 seconds relative to the peak of population-averaged pulsatile activity was used. Data from PPD 1–2, PPD 5–6, and PPD 11–12 were pooled and normalized, setting the peak activity for each ROI to 1. The resulting normalized data matrix (375 × 450) was subjected to principal component analysis (PCA) to reduce the dimensionality and extract the principal components (PCs) that capture the majority of the data variance. The PCA was performed using the MATLAB *pca* function, yielding PCs, scores, and other relevant outputs such as explained variance. We applied *k*-means clustering to the scores of the first six PCs, which accounted for more than 95% of the variance (Fig. S2). The *k*-means clustering was conducted with six clusters, using a fixed random seed for reproducibility.

To assign new samples to the existing clusters, we first normalized the new data using the mean obtained from the original 375 × 450 data matrix, ensuring that the new dataset was centered similarly to the original data. The centered new data were then projected into the PCA space using the PC vectors obtained from the original PCA. To assign the new samples to the predefined clusters, we used the *k*-nearest neighbors search to find the nearest cluster centroids from the original clustering using the MATLAB *knnsearch* function.

### Pup retrieval assay

Animals were placed in their home cage (191 × 376 × 163 mm) with standard wood chip bedding at least 1 day before the retrieval assay. Before the imaging session, the pups were removed, leaving two to three pups in the nest. After a 2–3-minute wait, the imaging session began. The retrieval assay was initiated by placing two pups in opposite corners of the nest. If the two pups were successfully retrieved, another two pups were placed in the same corners. This process continued until the 6-minute imaging session ended.

### Detection of pup-retrieval responsive cells in the PVH

To identify significant responsive cells during pup retrieval (Fig. 4), trial-averaged Ca^2+^ signals were compared between the behavioral events and a baseline period using the Mann–Whitney *U* test, with a significance threshold of p < 0.05. Behavior videos were recorded at 20 Hz using a camera (DMK 33UX174; Imaging Source). WaveSurfer generated precise transistor–transistor logic pulses to synchronize behavioral tracking with microendoscopic imaging. As described previously [48], cells were classified into one of six types, either elevated or suppressed in one of three behavioral categories. The following time windows were used for averaging: contact, 0 to +1 seconds after the initiation of contact; onset, 1.5 seconds after the onset of pup retrieval; and completion, 1.5 seconds after the completion of retrieval. The baseline time window was –1.5 to 0 seconds relative to each behavioral event. We first assigned elevated responding cells to contact, onset of retrieval, or completion, in this order. We noticed baseline fluctuations caused by suppressed responses in some cases. For example, a cell that showed a suppressed response during contact could be mistakenly assigned to an elevated responder during onset or completion because of a reduced baseline. In such cases, the baseline was recalculated from –1.5 to 0 seconds relative to the initiation of contact to reassess significance. Once all cells exhibiting elevated responses to one of the three behavioral categories were assigned, the remaining cells were classified if they showed significant suppressed responses to contact, onset of retrieval, or completion, in this order.

### Fiber photometry

Fiber photometry recordings were performed as previously described [20]. In brief, a 200 nL solution of AAV9 *Otp-GCaMP6s* was injected into the bilateral PVH of wild-type C57BL/6j female mice. Two weeks after the viral injection, a 400-μm core, N.A. 0.5 optical fiber (R-FOC-BL400C-50NA; RWD) was implanted approximately 100 μm above the PVH. After the surgery, the female mouse was housed singly except during mating or cohabitation with another mother and pups. Neurons expressing GCaMP were exposed to 465-nm light (modulated at 309.944 Hz) and 405-nm light (modulated at 208.616 Hz). The emitted fluorescence was captured using the integrated Fluorescence Mini Cube (iFMC4_AE (405)_E (460–490)_F (500–550)_S; Doric Lenses). Light acquisition, filtering, and demodulation were conducted with the Doric photometry setup and Doric Neuroscience Studio Software (Doric Lenses). The 405-nm signal was recorded as a background (non-calcium-dependent), while the 465-nm signal indicated the calcium-dependent GCaMP6s response. The power output at the fiber tip was approximately 5 μW, measured with a photodiode power sensor (S120VC; Thorlabs). The initial signal acquisition was set to 12 kHz and later decimated to 120 Hz. A 5-Hz low-pass filter was applied before analyzing the photometric peaks of OT neurons.

For the analyses, we used a custom Python code. Briefly, the background 405-nm signals were subtracted from the 465-nm signals after being fitted using the least-squares method. The ΔF/F (%) was then calculated as 100 × (F*t* – F_0_) / F_0_, where F_0_ was the average of the background-subtracted signals over the entire recording period and F*t* the background-subtracted signal at time *t*. Peak heights varied considerably among mothers, partly due to differences in optical fiber location relative to the PVH. To identify the photometric peaks reliably, we first selected several visually obvious peaks to estimate each animal’s peak height. The photometric peaks of the OT neurons were then automatically detected by using the *findpeaks* function in the Scipy module in Python, with a peak threshold set at half the estimated peak height and a FWHM threshold of over 1 second. To show the peri-event traces of the peaks, the peak was defined as the local maximum point of ΔF/F, with time 0 set as the point when the ΔF/F value reached half the peak height. ΔF/F data from –10 seconds to +15 seconds around time 0 were extracted, and the median fluorescence of the –8- to –3-second baseline period was adjusted to zero to align multiple data points along the y-axis.

### Histology and histochemistry

Mice were given an overdose of isoflurane and perfused transcardially with PBS followed by 4% paraformaldehyde (PFA) in PBS. Brain tissues were post-fixed with 4% PFA in PBS overnight at 4 °C, cryoprotected with 30% sucrose solution in PBS at 4 °C for 24–48 h, and embedded in O.C.T. compound (cat#4583; Tissue-Tek). For immunostaining of *GCaMP6s*, 40-μm coronal sections of the whole brain were collected using a cryostat (model #CM1860; Leica). Free-floating slices were incubated with gentle agitation at room temperature in the following solutions: 2 hours in blocking solution (5% heat-inactivated goat serum, 0.4% Triton-X100 in PBS); overnight at room temperature in primary antibody 1:1000 mouse anti-GFP (GFP-1010, Aves Labs) in blocking solution; 2–3 hours in secondary antibody 1:500 anti-chicken-IgY Alexa488-conjugated (Jackson ImmunoResearch cat# 703-545-155) in blocking solution; and 15 minutes in 2.5 μg/mL of DAPI (Santa Cruz, Cat #sc-3598) in PBS. Sections were mounted on slides and cover-slipped with mounting media (Fluoromount, Diagnostic BioSystems).

Sections were imaged using an Olympus BX53 microscope with a 4× (NA 0.16) or 10× (NA 0.4) objective lens equipped with a cooled CCD camera (DP80; Olympus) or a Zeiss Axio Scan.Z1 with a 10× (NA 0.45) objective lens.

### Quantification and statistics

For the analysis of microendoscopic data, statistical tests were performed using a custom MATLAB code. Photometry data analysis was performed with a custom Python code. All tests were two-tailed. The sample size and statistical tests used are detailed in the figures or figure legends. Statistical significance was set at P < 0.05. Data are presented as mean ± standard error of the mean (SEM), unless otherwise indicated.

### Data, Materials, and Software Availability

Custom MATLAB/Python codes supporting the findings of this study are available from the corresponding authors upon request. Data from fiber photometry and microendoscopy will be deposited in the SSDB depository (https://ssbd.riken.jp/repository/) by the time of publication.

## Supplementary Figures

**Figure S1.**
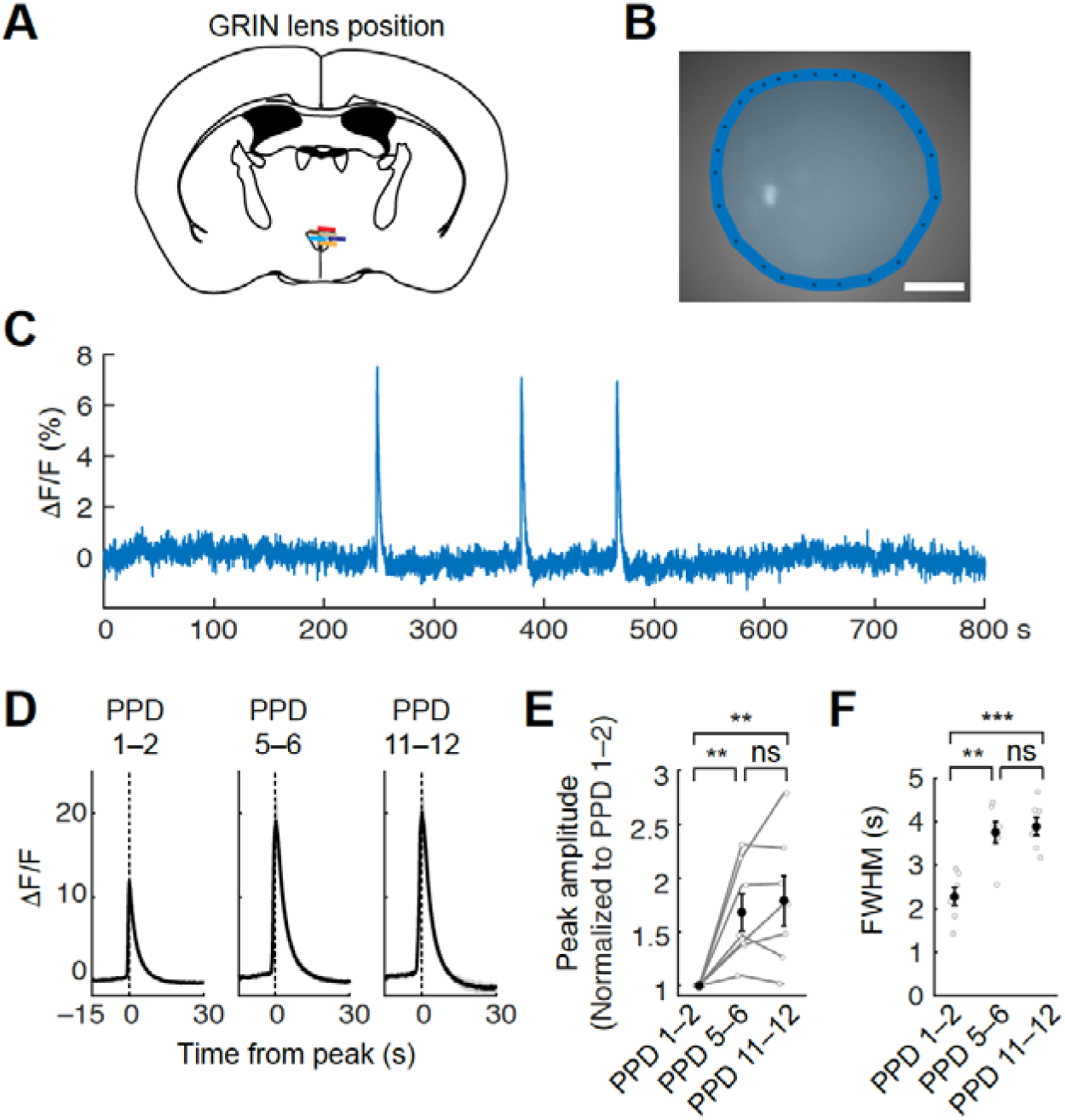
Measuring photometry-like population activity using microendoscopic imaging data, related to Fig. 2. (A) Schematics of coronal sections showing the position of the GRIN lens (N = 6 mice, one mouse was excluded due to death before sacrifice). (B) Example of a spatial map displaying a single ROI that covers the entire field of view of the GRIN lens. Scale bar, 100 μm. (C) Examples of Ca^2+^ responses from a single ROI in a PPD 1 mother during lactation. (D) Averaged activity traces during the pulsatile event. Peak time for each ROI was aligned to 0. N = 7 mice. Shadows represent SEM. (E, F) Quantification of the mean peak amplitude normalized to PPD 1–2 for each mouse (E) and FWHM (F). **, p < 0.01, and ***, p < 0.001 by post hoc Tukey’s HSD test after a significant Kruskal–Wallis test. Error bars, SEM.

**Figure S2.**
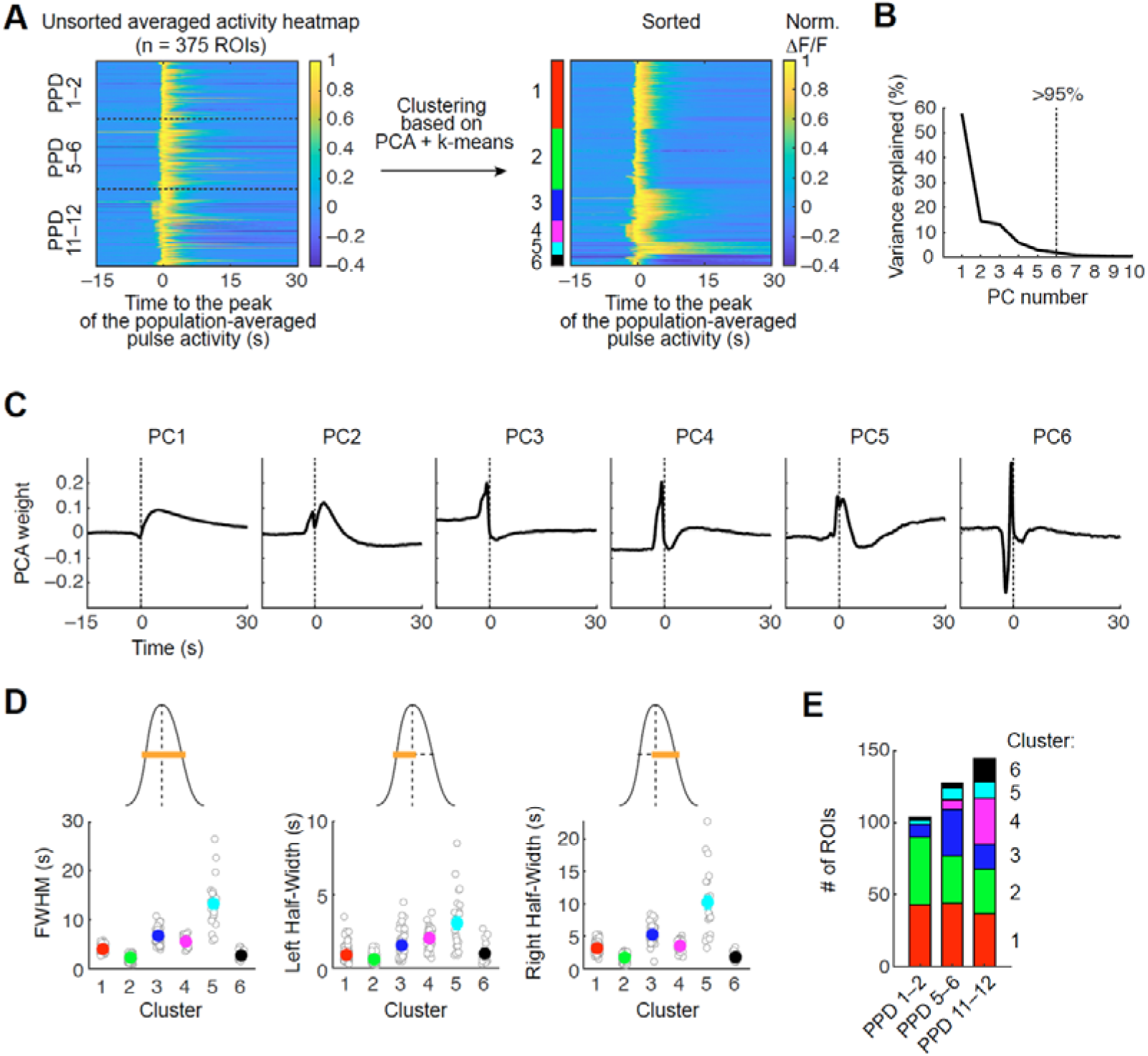
Clustering analysis, related to Fig. 2. (A) Normalized averaged response during pulsatile activity before (left) and after (right) clustering. (B) Scree plot of the percentage of explained variance per principal component. Over 95% of the variance was covered by up to six principal components (dashed line). (C) Individual retained principal components showing response vectors. (D) Quantifications of detailed parameters of waveforms in each cluster. (E) Number of ROIs classified into each cluster across different lactation stages.

**Figure S3.**
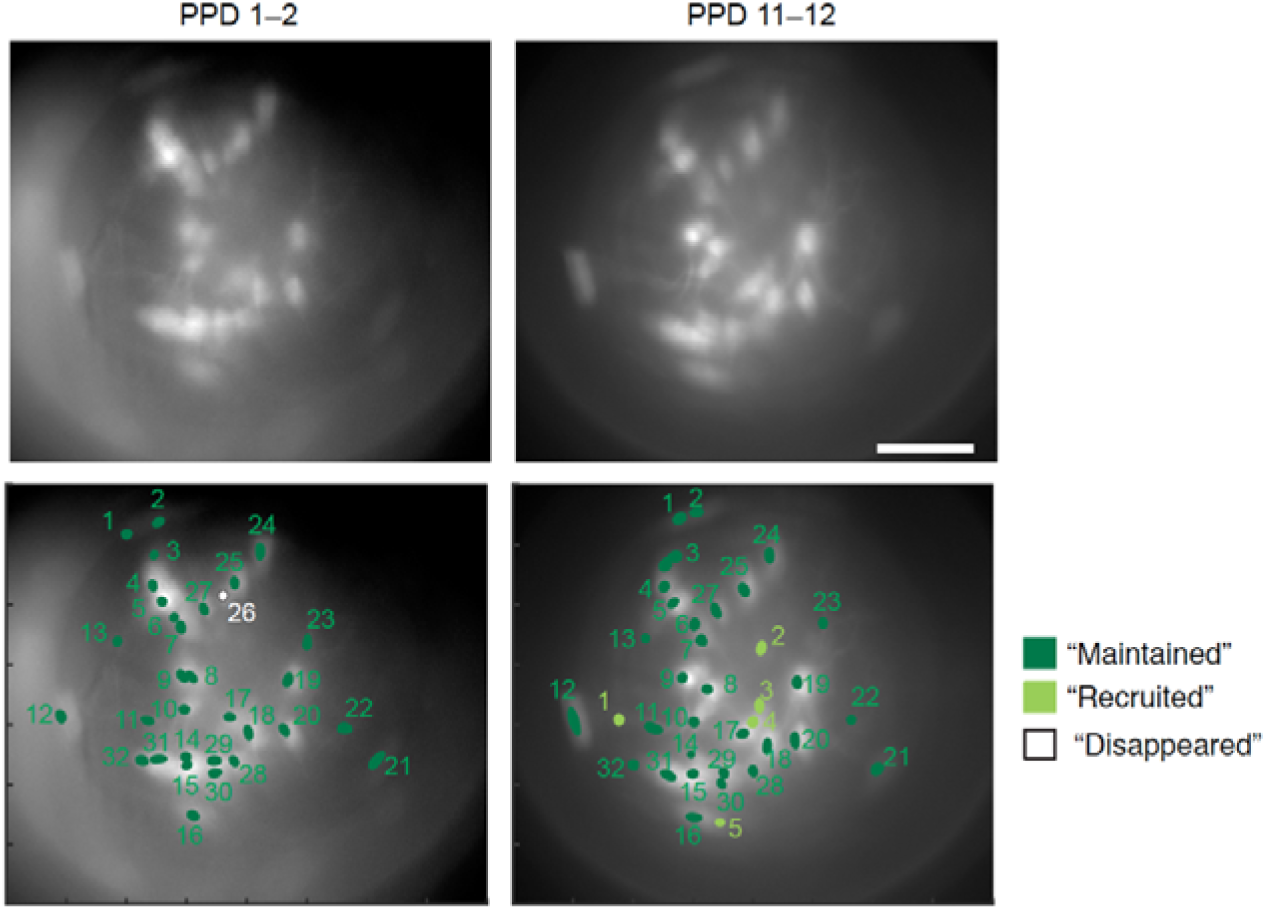
Longitudinal tracking, related to Fig. 3. “Maintained”, “Recruited”, and “Disappeared” ROIs from a representative animal. A total of 32 ROIs were detected at PPD 1–2, and 31 were reliably tracked at PPD 11–12 (dark green). Five ROIs were “Recruited” at PPD 11–12 (light green), and one ROI “disappeared”.

**Figure S4.**
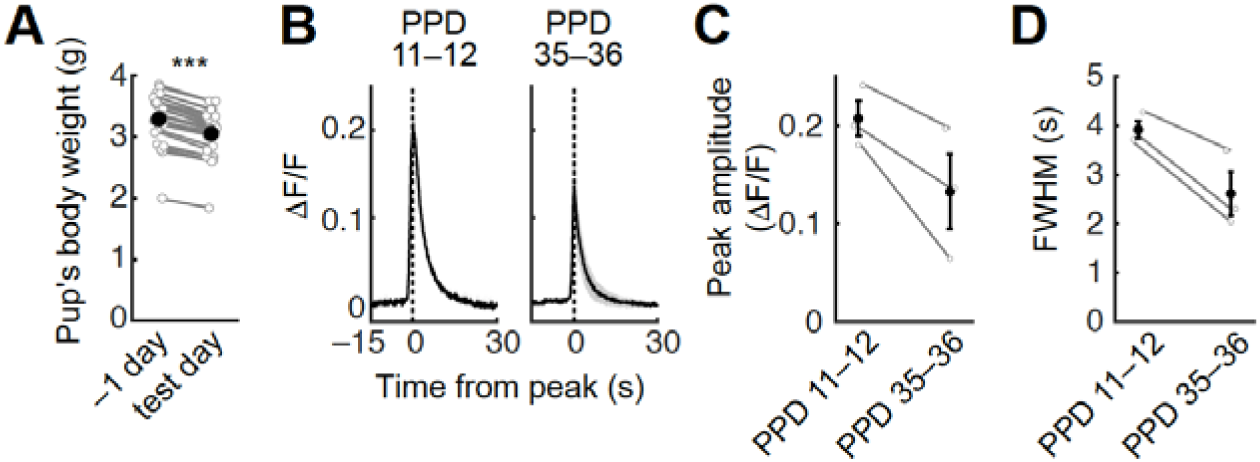
Photometry-like population activity of PVH OT neurons using microendoscopic imaging data, related to Fig. 5. (A) Quantification of the pup’s body weight after the imaging session and overnight co-housing with a post-weaning mother compared with the previous day. ***, p < 0.001 by a paired *t*-test. (B) Averaged activity traces during the pulsatile event. Peak time for each ROI was aligned to 0. N = 3 mice. Shadows represent SEM. (C, D) Quantification of the mean amplitude of the peak (C), and FWHM (D). Error bars, SEM.

**Figure S5.**
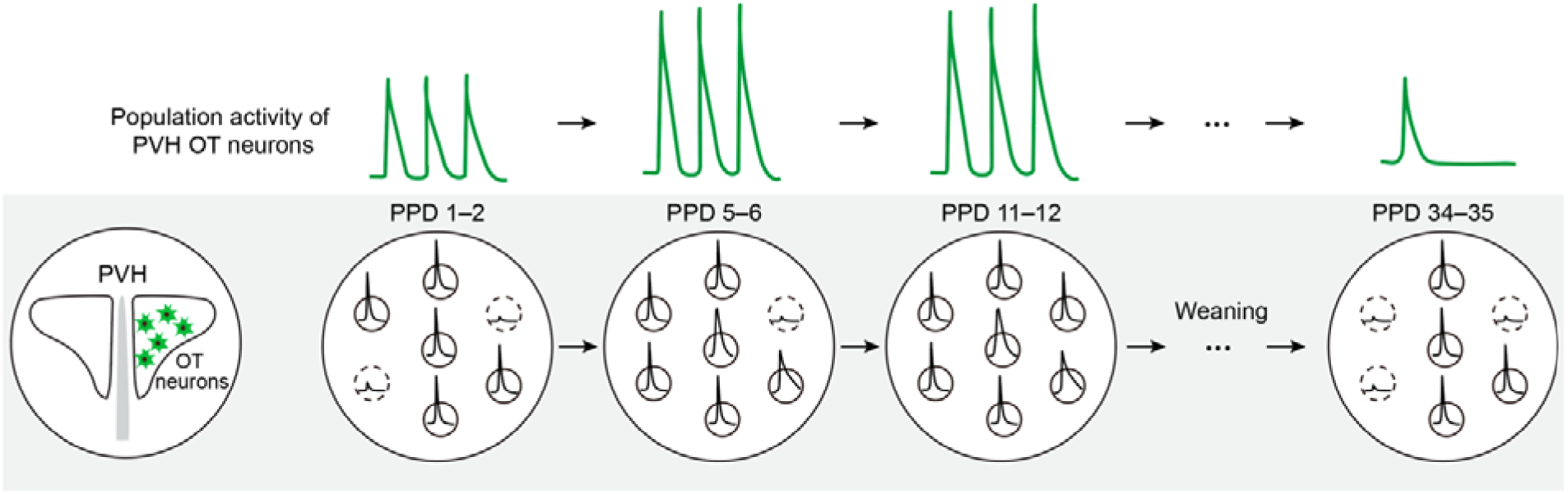
Graphical abstract. Top: Population pulsatile activity of PVH OT neurons induced by suckling stimuli. The amplitude increased during the mid-lactation stages, potentially enhancing milk ejection. Post-weaning mothers exhibited pulsatile activity with reduced amplitude. Bottom: Activity dynamics of individual PVH OT neurons. Changes in the population activity amplitude were primarily due to the recruitment and dismissal of responsive neurons, with waveforms of individual neurons also dynamically modulated across lactation stages.

